# CSF metabolites associate with CSF tau and improve prediction of Alzheimer’s disease status

**DOI:** 10.1101/2021.01.31.429054

**Authors:** Ruocheng Dong, Burcu F. Darst, Yuetiva Deming, Yue Ma, Qiongshi Lu, Henrik Zetterberg, Kaj Blennow, Cynthia M. Carlsson, Sterling C. Johnson, Sanjay Asthana, Corinne D. Engelman

## Abstract

**INTRODUCTION:** Cerebrospinal fluid (CSF) total tau (t-tau) and phosphorylated tau (p-tau) are biomarkers of Alzheimer’s disease (AD), yet much is unknown about AD-associated changes in tau metabolism and tau tangle etiology.

**METHODS:** We assessed the variation of t-tau and p-tau explained by 38 previously identified CSF metabolites using linear regression models in middle-age controls from the Wisconsin Alzheimer’s Disease Research Center, and predicted AD/mild cognitive impairment (MCI) vs. an independent set of older controls using metabolites selected by the least absolute shrinkage and selection operator (LASSO).

**RESULTS:** The 38 CSF metabolites explained 70.3% and 75.7% of the variance in t-tau and p-tau. Of these, 7 LASSO-selected metabolites improved the prediction ability of AD/MCI vs. older controls (AUC score increased from 0.92 to 0.97 and 0.78 to 0.93) compared to the base model.

**DISCUSSION:** These tau-correlated CSF metabolites increase AD/MCI prediction accuracy and may provide insight into tau tangle etiology.

## 1. Introduction

One of the defining neuropathological changes in Alzheimer’s disease (AD) is the intraneuronal aggregates of hyperphosphorylated and misfolded tau that give rise to neurofibrillary tangles and neuropil threads [1]. Their corresponding biomarkers in cerebrospinal fluid (CSF), total tau (t-tau) and phosphorylated tau (p-tau), can predict clinical AD and its progression [2]. Moreover, a new plasma p-tau biomarker (p-tau181) has recently been associated with AD pathology [3]. Research has been done to understand tau changes and how they happen [4,5]. For example, it has been shown that the dysregulation of kinases and phosphatases results in three to four times greater quantities of phosphorylated tau in the brains of AD patients than in normal adult brains [2], but the pathologic processes remain largely unknown.

Recent advancements in metabolomics technologies allow researchers to study multiple small molecules (<1500 Da), such as amino acids, fatty acids, and carbohydrates, simultaneously within a biological system [6]. Metabolites can be influenced by biological changes resulting from upstream molecular processes such as genetic mutations, as well as exogenous changes caused by environmental exposures (*e*.*g*., diet, medications, and physical activity). Moreover, compared to RNA transcripts and proteins, metabolites are more relevant to the current physiological state of a cell, and their abnormal levels and relative ratios can reflect disease progression, thus, metabolites serve as appropriate targets for health outcomes research [7].

To date, there have been numerous targeted or untargeted human blood metabolomic studies that focus on AD clinical status or CSF biomarkers [8]. For example, Toledo *et al*. [9] have conducted a network analysis using serum metabolites in participants from the Alzheimer’s Disease Neuroimaging Initiative and found that accumulation of acylcarnitine species indicates malfunction and alterations in tau metabolism. However, few studies have been conducted to assess the association between CSF metabolites and CSF tau. CSF communicates freely with the interstitial fluid that bathes the neurons and other cell types of the brain, spinal cord and the cranial and spinal nerves [10], which makes it an ideal source to study the pathological changes occurring in AD brains. By linking two well-established AD CSF biomarkers, CSF t-tau and p-tau, which reflect tau secretion and phosphorylation, and predict neurodegeneration and cortical tangle formation, respectively [11], with CSF metabolites, additional mechanistic information behind the development of pathological alterations related to tau may be revealed. The findings from studying CSF metabolites could ultimately be translated into potential AD prevention through modifiable risk factors (*e*.*g*., dietary interventions), better prognostic indicators, or new drug targets.

Darst *et al*. [12] constructed an inter-omics network consisting of whole blood gene expression, plasma metabolites, CSF metabolites, and AD risk factors in 1,111 non-Hispanic white participants from the Wisconsin Registry for Alzheimer’s Prevention (WRAP). Within this inter-omics network, a cluster of 38 CSF metabolites was identified in the subset of 141 individuals in which CSF was collected, with each individual metabolite being significantly correlated (p threshold: ≤6.1×10^−10^) with CSF t-tau and p-tau, and these collective metabolites accounting for 60.7% and 64.0% of the variation of t-tau and p-tau, respectively. In this study, we aimed to (1) replicate these findings and evaluate the predictive ability of these CSF metabolites in an independent sample (the IMPACT cohort) from the Wisconsin Alzheimer’s Disease Research Center (Wisconsin ADRC); (2) examine the predictive performance of the same metabolites present in plasma in WRAP; (3) identify the major metabolites driving this cluster in the IMPACT and WRAP cohorts and, in an independent sample, evaluate whether they can be used as potential biomarkers to enhance the prediction of AD or mild cognitive impairment (MCI); and (5) understand the biological functions of all 38 metabolites using pathway analyses to provide insight into disease-related processes. Our results confirm the previous associations between 38 CSF metabolites and CSF tau and provide potential biological mechanisms for the development of tau tangles and possible candidates for CSF metabolite biomarkers or drug targets.

## 2. Methods

### 2.1 Participants

The Wisconsin ADRC’s clinical core cohort started in 2009 and has well-characterized AD and MCI participants, as well as healthy older controls (HOC), and the IMPACT cohort of initially cognitively-unimpaired, asymptomatic middle-aged adults [13–15]. The replication sample for the main analysis included 158 non-Hispanic white individuals from the IMPACT cohort with cross-sectional CSF samples.

WRAP began recruitment in 2001 as a prospective cohort study of initially cognitively-unimpaired, asymptomatic, middle-aged adults enriched for a parental history of clinical AD [16]. The WRAP cohort included 130 and 123 non-Hispanic white individuals with longitudinal CSF and plasma samples, respectively. Both the CSF and plasma cohorts included five sibling pairs, one sibling trio and three sibling quartets. The WRAP dataset was utilized to reproduce and refine the results from Darst *et*.*al*. [12] using similar statistical models as those for the IMPACT cohort.

This study was conducted with the approval of the University of Wisconsin Institutional Review Board, and all participants provided signed informed consent before participation.

### 2.2 CSF and plasma sample collection and CSF biomarkers quantification

Fasting CSF samples for the Wisconsin ADRC cohorts and WRAP were collected via lumbar puncture [13] following the same protocol and by the same group of well-trained individuals [13]. Samples were sent together in two batches to the lab of Drs. Blennow and Zetterberg in Sweden, where commercially available enzyme-linked immunosorbent assay (ELISA) methods were used to quantify CSF t-tau, p-tau, and amyloid-beta 1-42 (Aβ_42_) (INNOTEST® assays HTAU AG, PHOSPHO-TAU[181P], and β-amyloid1-42, respectively; Fujirebio, Ghent, Belgium) [13]. The batch-adjusted predicted values for CSF biomarkers were used for all analyses [17].

In WRAP, fasting blood samples were collected in ethylenediaminetetraacetic acid (EDTA) tubes; the plasma was pipetted off within one hour of collection and stored at -80°C [12]. A total of 141 longitudinal samples from 123 individuals in WRAP with plasma metabolites were available for the main analysis. In the Wisconsin ADRC, blood samples were collected in heparin tubes, which could influence metabolite values; as such, plasma metabolomics data have not been generated in Wisconsin ADRC blood samples. Further details of how plasma and CSF samples were processed are explained in an earlier study [12].

### 2.3 CSF metabolomic profiling and quality control

CSF and plasma metabolomic analyses and quantification were performed in one batch by Metabolon (Durham, NC) using an untargeted approach, based on Ultrahigh Performance Liquid Chromatography-Tandem Mass Spectrometry platform (UPLC-MS/MS) [18]. Details of the metabolomic profiling were described in an earlier study [12].

Each metabolite value was first scaled so the median was equal to one across all samples. Missing values were then imputed to half the lowest level of detection for each biological metabolite and 0.0001 (the lowest value that could be accepted in the analytic software) for each xenobiotic metabolite. The missing percentage for each of the 38 previously identified CSF metabolites prior to imputation is shown in Supplemental Table 1. Metabolites with zero variability between individuals, or with an interquartile range of zero, were excluded (none of the 38 CSF metabolites were excluded). Log10 transformation was employed to normalize the data. After quality control, the previously identified 38 metabolites were selected for this investigation. The distribution of each of the 38 CSF metabolites after imputation and Log 10 transformation is shown in Supplemental Figure 1.

### 2.4 Statistical analysis

#### 2.4.1 Prediction performance of the 38 CSF metabolites

To replicate the previously reported results in WRAP [12], each metabolite’s association with t-tau and p-tau was tested in the IMPACT cohort and the Bonferroni adjustment was applied to correct for multiple testing. A meta-analysis was also conducted by using results from IMPACT and WRAP. To replicate the performance of the cluster of 38 CSF metabolites in explaining variation in tau pathology, we used linear regression models to determine the prediction performance (r^2^) of CSF t-tau and p-tau in IMPACT. The base models, which included age, sex, and years of education, were compared to models that also included the 38 CSF metabolites. To reproduce the results in WRAP and compare them to IMPACT using consistent statistical models, we determined the prediction performance (r^2^) of the 38 CSF metabolites using linear mixed-effects regression with random intercepts to account for repeated measures and sibling relationships. In both IMPACT and WRAP, we randomly split the data into a training (70%) and validation (30%) set and created plots to compare the observed and predicted values. Finally, we physically combined the WRAP baseline samples and IMPACT samples and re-conducted the analysis to evaluate the explained variance. Sex-stratified prediction differences were assessed in WRAP by fitting the mentioned models in males and females separately. The number of male samples in IMPACT was too small to perform sex-stratified analyses while meeting the degrees of freedom needed by the model. Of the original 38 CSF metabolites, 34 were also found in plasma samples from WRAP and were tested together as predictors for t-tau and p-tau using linear mixed-effects regression models, as described above. A sensitivity analysis using only the baseline samples was also conducted in WRAP for both CSF and plasma metabolites. The statistical analyses here and below were all conducted in R version 3.6.2. The lme4 package was used.

#### 2.4.2 LASSO selection of important metabolites and their prediction of AD/MCI vs. HOC

In order to incorporate a practical number of metabolites in the prediction model of AD/MCI diagnosis vs. HOC instead of including all 38 metabolites, the least absolute shrinkage and selection operator (LASSO) [19] was applied to select the most important metabolites (those with non-zero estimated effects) for CSF t-tau and p-tau in both IMPACT and WRAP. In WRAP, the average of longitudinal CSF measures was used in LASSO regression. The tau variances explained by the selected metabolites were re-evaluated using similar model from 2.4.1 in both IMPACT and WRAP. The ability to enhance the prediction of AD/MCI vs. HOC status by the metabolites selected from LASSO was evaluated in an independent set of participants from the Wisconsin ADRC using logistic regression and an area under the curve (AUC) score. To determine prediction ability of the selected metabolites beyond demographic factors and established biomarkers, base models including age, sex, years of education, *APOE* ε4 count, t-tau, p-tau, and Aβ_42_ were compared to the base model replacing t-tau and p-tau with the selected metabolites and also the base model plus the selected metabolites from LASSO. The analysis here used the “glmnet” package in R.

#### 2.4.3 Biological relevance of the 38 CSF metabolites

An exploratory factor analysis was conducted to determine if subsets of metabolites clustered together in latent factors associated with t-tau and p-tau. The factor analysis was performed in IMPACT and WRAP using the “psych” package in R. Metabolites with a loading of ≥0.4 [20] in one particular factor and lower loadings for the rest of the factors were considered as members of that particular factor. Potential functional pathways of the 38 metabolites were identified from the Homo sapiens KEGG pathway by conducting pathway analyses using the web-based software, Metabo-analyst [21], inputting the metabolites’ human metabolome database (HMDB) IDs, and using the default hypergeometric test and the relative-betweenness centrality, which is a measure of centrality in a graph based on the shortest paths that pass through the vertex. Pathways were considered as important if the FDR was ≤0.05 or the impact was ≥0.1.

## 3. Results

### 3.1 Participant characteristics

Characteristics of the participants can be found in Table 1. Among 158 Wisconsin ADRC IMPACT participants and 130 WRAP participants who had CSF metabolite data available, females comprised 74.7% of IMPACT participants and 65.4% of participants in WRAP. The mean baseline age was significantly younger in IMPACT (57.8 years) compared to WRAP (61.5 years). The mean years of education was similar (16.0 and 16.1 years in IMPACT and WRAP, respectively). Mean CSF t-tau was significantly lower in IMPACT (283.1) compared to WRAP (311.5). The correlation between t-tau and p-tau was approximately 0.90 in IMPACT and WRAP. The characteristics of each additional sub cohort of the Wisconsin ADRC and of the 123 WRAP participants in the plasma prediction analysis can also be found in Table 1.

**Table 1.** Wisconsin ADRC and WRAP participant characteristics.

| Participant Characteristics | CSF |  |  |  |  |  | Plasma |
| --- | --- | --- | --- | --- | --- | --- | --- |
|  | Wisconsin ADRC |  |  |  | WRAP<br>(n = 130<sup>†</sup>) | P<br>values* | WRAP<br>(n = 123) |
|  | AD<br>(n = 38) | MCI<br>(n = 29) | HOC<br>(n = 40) | IMPACT<br>(n = 158) |  |  |  |
| **Female:** n (%) | 12 (31.6) | 8 (27.6) | 22 (55.0) | 118 (74.7) | 85 (65.4) | 0.11 | 80 (65.0) |
| **Age in years:**<br>mean (SD) | 71.4 (8.9) | 74.2 (8.3) | 74.1 (4.8) | 57.8 (5.3) | 61.5 (6.6) | <0.001 | 62.6 (6.5) |
| **Education in years:**<br>mean (SD) | 14.8 (2.9) | 16.3 (2.8) | 16.4 (2.9) | 16.0 (2.3) | 16.1 (2.2) | 0.50 | 16.1 (2.2) |
| **CSF t-tau:**<br>mean (SD) | 775.5<br>(332.9) | 573.0<br>(346.9) | 436.8(194.5) | 283.1<br>(144.2) | 311.5<br>(117.7) | 0.03 | 318.9<br>(122.7) |
| **CSF p-tau:**<br>mean (SD) | 74.8<br>(27.3) | 62.4(36.2) | 52.9 (19.4) | 39.8<br>(14.5) | 41.5<br>(13.4) | 0.10 | 42.6<br>(13.5) |

\* The p values are based on a two-sample t test conducted between the Wisconsin ADRC IMPACT cohort and WRAP (shaded columns). <sup>†</sup>The 130 individuals had 210 longitudinal samples.

### 3.2 Prediction performance

Each of the 38 CSF metabolites was significantly associated with t-tau and p-tau in IMPACT and the direction of the effect was the same as in WRAP (Supplemental Table 2). Meta-analysis results are shown in Supplemental Table 3. All metabolites were significantly associated with t-tau and p-tau except erythritol. Base models only explained approximately 10% of the variance in t-tau and p-tau in both IMPACT and WRAP (Table 2). In IMPACT, the statistical model including the 38 CSF metabolites and demographics together explained 70.3% of the variance in t-tau and 75.7% of the variance in p-tau values. These results were similar to those calculated in WRAP, where the model including the 38 CSF metabolites and demographics explained 62.4% and 65.1% of the variance in t-tau and p-tau values, respectively. Similarly, in the combined dataset, the 38 CSF metabolites explained 66.1% and 72.3% of the variance in t-tau and p-tau, respectively. The results of the same analysis but only using baseline samples in WRAP are shown in Supplemental Table 4. Supplemental Figure 2 shows plots comparing the observed and predicted values for t-tau and p-tau in both IMPACT and WRAP. In WRAP, these metabolites explained more of the variance in the t-tau and p-tau in males (r^2^=0.749 and 0.804) than in females (r^2^=0.591 and 0.640; Table 2). We did not have enough male participants to fit the sex-stratified model in IMPACT; however, while the female only r^2^ was lower than the overall r^2^ in WRAP, this trend was not seen in IMPACT. In WRAP, the 34 of 38 metabolites present in plasma explained 26.9% and 30.1% of the variance in CSF t-tau and p-tau, respectively (Table 2), which is relatively low compared to CSF metabolites. We also examined the same 34 CSF metabolites’ prediction ability and confirmed that the lower r^2^ values for the 34 plasma metabolites were not due to the absence of the four metabolites (Supplemental Table 4).

**Table 2.** Prediction performance (r^2^) of each model in IMPACT and WRAP.

|  | IMPACT |  | WRAP |  |
| --- | --- | --- | --- | --- |
|  | T-tau | P-tau | T-tau | P-tau |
| **Null models: demographics\* only** | 0.083 | 0.106 | 0.085 | 0.087 |
| **Enhanced models: demographics and 38 CSF metabolites** |  |  |  |  |
| Overall | 0.703 | 0.757 | 0.624 | 0.651 |
| Female | 0.787 | 0.791 | 0.591 | 0.640 |
| Male | NA† | NA† | 0.794 | 0.804 |
| **Demographics and 7 LASSO selected CSF metabolites** | 0.594 | 0.692 | 0.585 | 0.615 |
| **Demographics and 34 Plasma metabolites** | NA† | NA† | 0.269 | 0.301 |

\* Demographics are age, sex, and years of education.
† The sample size for males was too small to perform the analysis in IMPACT.

### 3.3 LASSO results

LASSO results for t-tau and p-tau in both IMPACT and WRAP are shown in Table 3. Eight metabolites with non-zero coefficients (ranging from 33.25 to 202.10) were chosen in IMPACT, and twelve metabolites (coefficients ranging from - 112.48 to 333.57) were selected for t-tau in WRAP. Among the selected metabolites, five were consistent across IMPACT and WRAP (N-acetylneuraminate, C-glycosyl tryptophan, X-10457, X-24228, and 1-palmitoyl-GPC(16:0)). Eleven metabolites in IMPACT and twelve metabolites in WRAP with non-zero coefficients (ranging from 1.07 to 28.80 in IMPACT and -4.06 to 30.80 in WRAP) were selected for p-tau, with seven metabolites overlapping (N-acetylneuraminate, C-glycosyl tryptophan, X-10457, X-24228, 1-oleoyl-GPC(18:1), 1-palmitoyl-GPC(16:0), and 1-myristoyl-2-palmitoyl-GPC(14:0/16:0)), which included the five metabolites overlapping in the two t-tau models. These seven metabolites along with demographics explained about 59% and 69% of the variance in t-tau and p-tau, respectively in IMPACT and 59% and 62%, respectively in WRAP (Table 2).

**Table 3.** LASSO results for CSF t-tau and p-tau in IMPACT and WRAP.

| IMPACT |  | WRAP |  |
| --- | --- | --- | --- |
| T-tau |  |  |  |
| Biochemical Name | Coefficient | Biochemical Name | Coefficient |
| X - 24228 | 202.10 | N-acetylneuraminate | 333.57 |
| N-acetylneuraminate | 188.68 | C-glycosyl tryptophan | 264.20 |
| Beta-citrylglutamate | 151.01 | X - 12906 | 97.42 |
| C-glycosyl tryptophan | 92.25 | 1-oleoyl-GPC (18:1) | 69.00 |
| N-acetylthreonine | 78.41 | X - 10457 | 62.67 |
| 1-palmitoyl-GPC (16:0) | 72.78 | N6-succinyladenosine | 50.65 |
| X - 10457 | 42.70 | X - 24228 | 36.65 |
| X - 24699 | 33.25 | X - 23739 | 28.95 |
|  |  | 1-palmitoyl-GPC (16:0) | 1.38 |
|  |  | 1-palmitoyl-2-palmitoleoyl-GPC (16:0/16:1) | 0.21 |
|  |  | X - 18887 | -3.56 |
|  |  | Ribonate | -112.48 |
| P-tau |  |  |  |
| C-glycosyl tryptophan | 28.80 | N-acetylneuraminate | 30.80 |
| N-acetylneuraminate | 24.98 | C-glycosyl tryptophan | 30.55 |
| Beta-citrylglutamate | 15.71 | X - 10457 | 9.80 |
| X - 24228 | 7.37 | N6-succinyladenosine | 7.41 |
| X - 10457 | 5.47 | X - 24228 | 5.59 |
| 1-palmitoyl-GPC (16:0) | 4.26 | X - 12906 | 3.66 |
| Cholesterol | 2.52 | Sphingomyelin (d18:1/14:0, d16:1/16:0) | 3.05 |
| 1-oleoyl-GPC (18:1) | 2.44 | X - 24699 | 3.04 |
| X - 24329 | 1.25 | 1-oleoyl-GPC (18:1) | 2.33 |
| Gulonate | 1.13 | 1-myristoyl-2-palmitoyl-GPC (14:0/16:0) | 0.88 |
| 1-myristoyl-2-palmitoyl-GPC (14:0/16:0) | 1.07 | X - 18887 | -2.50 |
|  |  | Ribonate | -4.06 |

Metabolites shaded in light grey with bold font are consistent across IMPACT and WRAP.

When predicting AD vs. HOC and MCI vs. HOC, the base models, including age, sex, years of education, *APOE* ε4 count, t-tau, p-tau, and Aβ_42_, achieved AUC scores of 0.92 and 0.78, respectively. Replacing t-tau and p-tau with the seven metabolites selected by LASSO, that overlapped across IMPACT and WRAP for t-tau and/or p-tau, achieved AUC scores of 0.94 and 0.82. The base model plus the seven metabolites collectively improved the prediction ability of AD vs. HOC (AUC score increased from 0.92 to 0.97) and of MCI vs. HOC (AUC score increased from 0.78 to 0.93; Figure 1). The comparisons of results from the base model plus seven LASSO selected metabolites to the base model with seven randomly selected metabolites from 38 metabolites and seven randomly selected metabolites from all CSF metabolites with tau outcomes are shown in Supplemental Figure 3.

**Figure 1.**
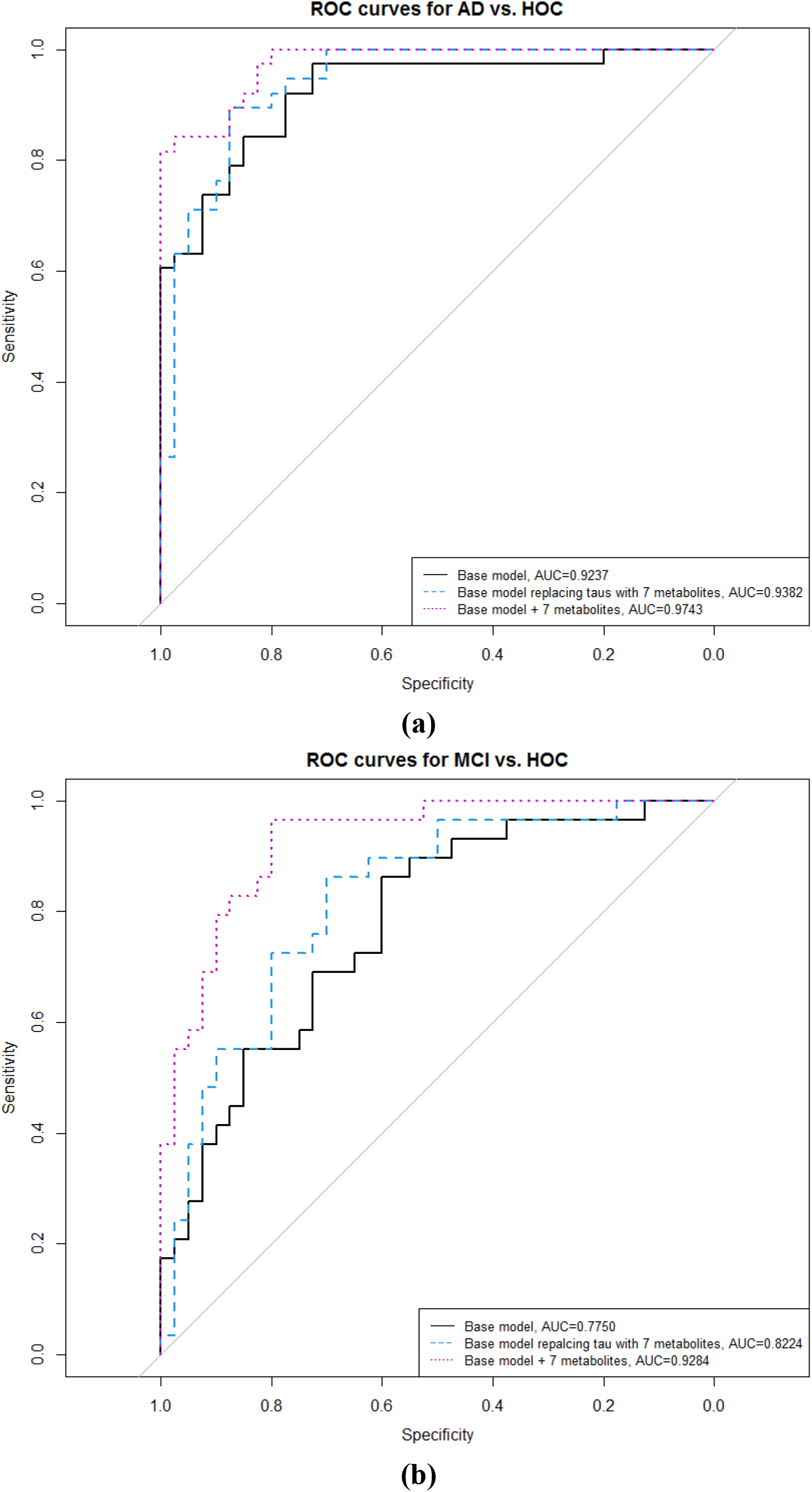
ROC curves and AUC scores of predictions by 6 models in the Wisconsin ADRC (a) AD vs. HOC (b) MCI vs. HOC. Base model: age, sex, years of education, *APOE* ε4 count, t-tau, p-tau, and Aβ42; base model replacing t-tau and p-tau with the seven selected metabolites from LASSO; and base model plus the seven selected metabolites from LASSO.

### 3.4 Biological relevance of the 38 metabolites

The biochemical names, sub-pathways, and super pathways of the 38 metabolites can be found in Supplemental Table 5, which also shows the loadings of each metabolite for three latent factors produced through exploratory factor analysis. These factors included the exact same metabolites and similar loadings for each in both IMPACT and WRAP and explained about 60% of the variance in the 38 metabolites. Factor 1 included 25 metabolites in the following pathways: amino acids, nucleotides, carbohydrates, cofactors and vitamins, energy, xenobiotics, and unknowns (no confirmed biochemical names). Factor 2 was composed of eleven lipids. Two lysophospholipids contributed to factor 3 and they were selected by LASSO for p-tau in both IMPACT and WRAP.

Among the 29 known metabolites, 26 had HMDB IDs and 23 of these were present in the MetaboAnalyst database. In pathway analyses, these 23 metabolites were enriched in two KEGG pathways (Figure 2 and Table 4): (1) pentose and glucuronate interconversions and (2) glycerophospholipid (GP) metabolism. Three metabolites from Factor 1, arabinose, arabitol/xylitol, and gulonate, were enriched in pentose and glucuronate interconversions. Two metabolites, 1-palmitoyl-2-palmitoleoyl-GPC(16:0/16:1) and 1-oleoyl-GPC(18:1) from Factors 2 and 3, respectively, were enriched in glycerophospholipid metabolism.

**Table 4.** Pathway analysis results for the 38 CSF metabolites.

|  | Pathway Enrichment |  |  |  |  | Pathway Impact |
| --- | --- | --- | --- | --- | --- | --- |
| KEGG Pathway | Total # Metabolites in KEGG Pathway | # Metabolites Identified in Present Study | Raw p | -Log(p) | FDR | Impact |
| Pentose and glucuronate interconversions | 18 | 3 | 2E-04 | 8.80 | 0.01 | 0.25 |
| Glycerophospholipid metabolism | 36 | 2 | 0.02 | 3.86 | 0.88 | 0.11 |
| Linoleic acid metabolism | 5 | 1 | 0.03 | 3.45 | 0.89 | 0 |
| Ascorbate and aldarate metabolism | 8 | 1 | 0.05 | 2.98 | 1 | 0 |
| alpha-Linolenic acid metabolism | 13 | 1 | 0.08 | 2.51 | 1 | 0 |
| Sphingolipid metabolism | 21 | 1 | 0.13 | 2.06 | 1 | 0 |
| Arachidonic acid metabolism | 36 | 1 | 0.21 | 1.56 | 1 | 0 |
| Steroid biosynthesis | 42 | 1 | 0.24 | 1.42 | 1 | 0.03 |
| Primary bile acid biosynthesis | 46 | 1 | 0.26 | 1.34 | 1 | 0.05 |
| Steroid hormone biosynthesis | 85 | 1 | 0.43 | 0.84 | 1 | 0.005 |

Raw p is the original p value calculated from the enrichment analysis; FDR p is the p value adjusted using False Discovery Rate; Impact is the pathway impact value calculated from the pathway topology analysis. The shaded two pathways are considered important.

**Figure 2.**
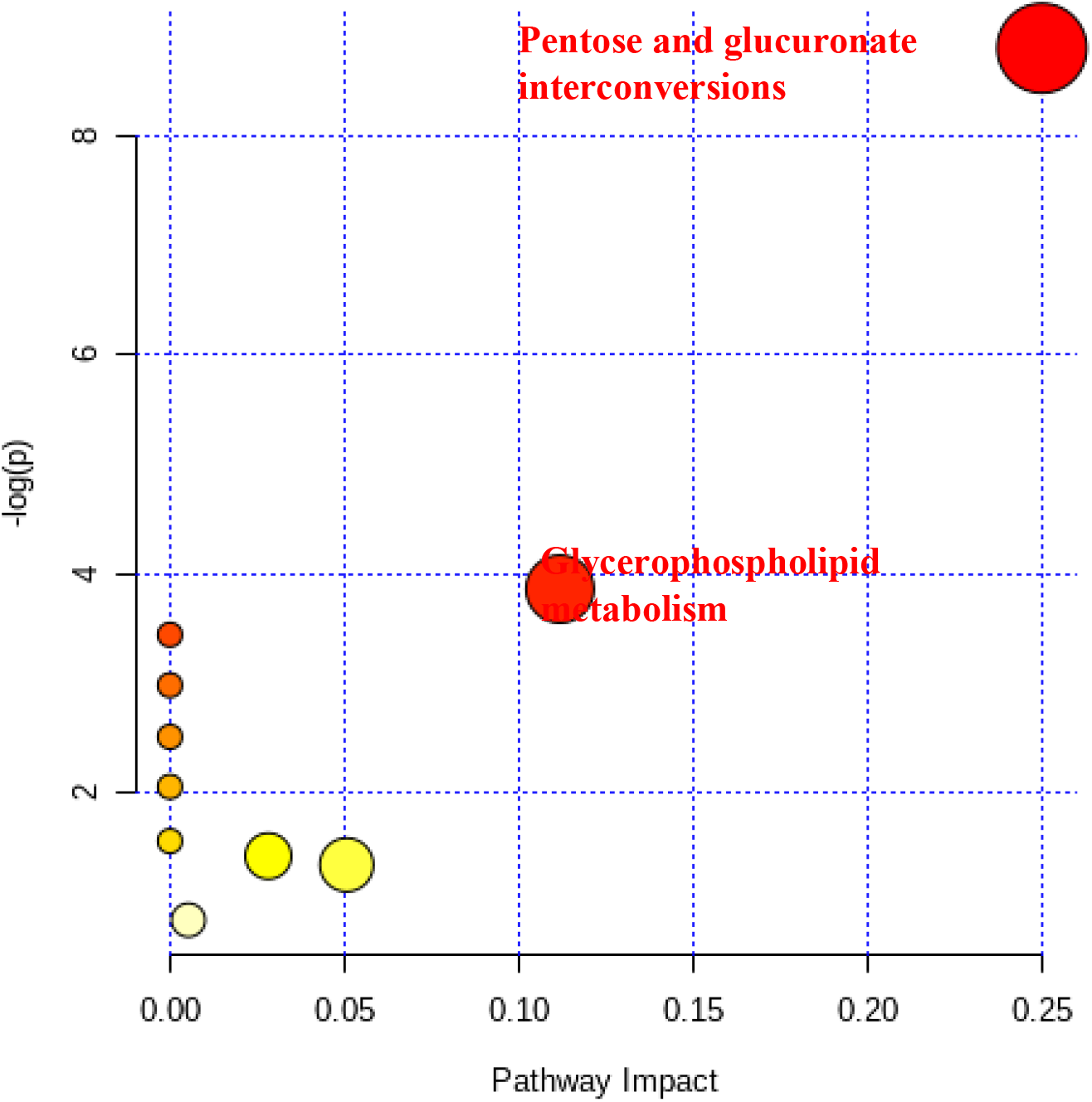
Pathway analysis results for 23 CSF metabolites. The x-axis represents the pathway impact, and y-axis represents the pathway enrichment. Larger sizes and darker colors represent higher pathway impact and enrichment, respectively.

## 4. Discussion

Using a cross-sectional sample from the Wisconsin ADRC IMPACT cohort, we replicated previous findings of 38 CSF metabolites associated with t-tau and p-tau in WRAP [12]. Not only was each of the 38 CSF metabolites significantly associated with both tau outcomes after Bonferroni correction, but the high amount of variance in tau explained by this cluster of 38 CSF metabolites was confirmed in IMPACT.

Among these metabolites, there are 13 lipids, 7 amino acids, 5 carbohydrates, 1 nucleotide, 1 energy metabolite, 1 cofactor and vitamin metabolite, 1 xenobiotic, and 9 unknown metabolites. Some of these metabolites, such as 1,2-dipalmitoyl-GPC(16:0/16:0) and stearoyl sphingomyelin(d18:1/18:0), were previously reported to be associated with AD diagnosis or AD pathogenesis [22,23]. Orešič et al. (2011) found that serum 1,2-dipalmitoyl-GPC(16:0/16:0), also called PC(16:0/16:0), was one of 3 metabolites considered to be predictive markers of AD progression in individuals with MCI [22]. CSF stearoyl sphingomyelin(d18:1/18:0), also called SM(d18:1/18:0), distinguished clinical AD from controls, with an accuracy of 70% and was significantly increased in patients displaying pathological levels of Aβ_42_, t-tau and p-tau [23], supporting that this molecule changes in patients with A/T/N pathology. Additionally, the N-acetylamino acids, N-acetylvaline, N-acetylthreonine, N-acetylserine, and N-acetyl-isoputreanine, were identified in our study. N-acetylthreonine and N-acetylserine are the downstream metabolites of the cleavage process initiated by lysosomal protease tripeptidyl peptidase 1 (TPP1) [24], and previous studies [25,26] suggested that increased levels of TPP1 enhance fibrillar β-amyloid degradation. In support of this, a secondary analysis in our study found that N-acetylserine was significantly associated with Aβ_42_ (beta=480.38, p=0.002), providing evidence that this CSF metabolite may be involved in brain amyloid pathology.

From the 38 metabolites, 7 were selected by LASSO in both IMPACT and WRAP: N-acetylneuraminate, C-glycosyl tryptophan, 1-palmitoyl-GPC(16:0), 1-oleoyl-GPC(18:1), 1-myristoyl-2-palmitoyl-GPC(14:0/16:0), and two unknown metabolites (X-10457 and X-24228). These improved the prediction of AD vs. HOC by approximately 5% and MCI vs HOC by 15% compared to a model that included the well-established AD risk factors of age, sex, years of education, *APOE* ε4 count, t-tau, p-tau, and Aβ_42_. A recent study in a Japanese cohort found that CSF N-acetylneuraminate was significantly higher in AD patients, when compared to the idiopathic normal pressure hydrocephalus, and had a positive correlation with CSF p-tau (r=0.55) [27]. In our study, CSF N-acetylneuraminate was positively associated with both t-tau and p-tau. C-glycosyl tryptophan, a sugar-loaded amino acid, has been reported to be strongly associated with aging, defined by chronological age (beta=2.47, p=1.3×10^−23^), in a human blood metabolome-wide association study [28]. In our study, CSF C-glycosyl tryptophan was positively associated with t-tau and p-tau. Two lipids 1-palmitoyl-GPC(16:0) (also called LysoPC(16:0/0:0)) and 1-myristoyl-2-palmitoyl-GPC(14:0/16:0) (also called PC(14:0/16:0)), belong to the class of lysophospholipid (LysoPCs) and phosphatidylcholines (PCs), respectively. Previous studies have shown that numerous plasma/serum metabolites from the LysoPC and PC classes were significantly associated with MCI and AD dementia or able to discriminate MCI and AD dementia cases from controls [9,29–34]. In a randomized crossover trial that treated mild to moderate AD patients with medium-chain triglycerides, 1-palmitoyl-GPC(16:0) levels increased along with an improvement in cognition [34]. These seven LASSO-selected metabolites improved the prediction of AD and MCI status, suggesting they may be useful biomarkers for clinical AD and MCI diagnosis.

Another interesting discovery from the LASSO results is that, while five metabolites were overlapping in IMPACT and WRAP for both t-tau and p-tau, two metabolites,1-oleoyl-GPC(18:1) (also called LysoPC(18:1(9Z)/0:0)) and 1-myristoyl-2-palmitoyl-GPC(14:0/16:0) (also called PC(14:0/16:0)) were selected only for p-tau, not t-tau. Since p-tau is more specific to AD-related tau pathology than t-tau, these metabolites might provide insight into the pathological processes involved in tau tangle formation in AD.

When using the seven metabolites to predict AD/MCI vs. HOC, the AUC scores from both the base model and base model+metabolites were higher for AD than for MCI. However, a greater improvement in prediction accuracy for MCI vs. HOC (15%) was achieved than AD vs. HOC (5%). One possible reason could be that the 38 CSF metabolites were originally identified from the WRAP cohort, whose participants were relatively young and have not been diagnosed with AD yet. Another explanation could be that the base model, which included demographics, APOE ε4 count, and three core AD CSF biomarkers, already achieved a very high accuracy for predicting AD vs. HOC and had little room for improvement.

In WRAP, we were able to test the prediction of 34 of the 38 metabolites that were found in plasma. We found that these 34 metabolites collectively did not explain much variation in CSF concentrations of t-tau and p-tau (r^2^ between 0.286 and 0.303). This was not due to the absence of the 4 metabolites, since the r^2^ of the 34 metabolites in the CSF (0.621 to 0.641) was close to that with all 38 metabolites (0.624 to 0.651) in WRAP. Moreover, the correlations between the same 34 metabolites measured in both CSF and plasma are relatively low (−0.13 to 0.30) [12] (Supplemental Figure 4). We previously proposed that this low correlation could be attributed to these metabolites not being able to cross the blood brain barrier (BBB) [35]. For example, cholesterol metabolism in the brain relies on its own cells to produce cholesterol, and the transport of cholesterol from peripheral circulation into the brain is prevented by the BBB [36,37]. In this situation, the concentrations and functions of metabolites like cholesterol are different across the BBB. Thus, testing for these metabolites in a more readily available body fluid, like blood, does not appear to be a viable option.

The factor analysis results suggest that the 38 metabolites are associated with tau through three main clusters (1) the combination of select amino acids, nucleotides, carbohydrates, cofactors and vitamins, energy, xenobiotics, and unknown metabolites; (2) phosphatidylcholines and sphingolipid metabolism, and (3) lysophospholipids. Five metabolites from these factors were enriched in (1) pentose and glucuronate interconversions and (2) glycerophospholipid metabolism from the pathway analysis. The pentose and glucuronate interconversion pathway was suggested from genomics and metabolomics studies to be involved in AD [38–40]. A urinary metabolomics study of APP/PS1 transgenic mice of AD and a hippocampal metabolomics study of CRND8 mice also identified this pathway [41,42]. Other studies have shown that brain glucose dysregulation and pentose related activities are associated with AD pathology [43–46]. Thus, our results provide further potential links between molecules in pentose and glucuronate metabolism, especially the three CSF metabolites, arabinose, xylitol, and gulonate, and the tau pathological process of AD.

The brain is the most cholesterol-rich organ, containing glycerophospholipids, cholesterol, sphingolipids, etc. [47]. The neural membranes are also composed of these lipids and the evidence suggests that glycerophospholipids and glycerophospholipid metabolism may associate with neural membrane composition alterations, glycerolipid-derived lipid-mediated oxidative stress, and neuroinflammation [9,48]. For example, levels of glycerophospholipids were decreased in brain autopsy samples from AD patients compared to age-matched controls [49]. In another study, increased glycerophospholipid levels were associated with increased activities of lipolytic enzymes and elevated concentrations of phospholipid degradation metabolites [50]. In our analysis, the two metabolites, 1-palmitoyl-2-palmitoleoyl-GPC(16:0/16:1) and 1-oleoyl-GPC(18:1), from Factors 2 and 3 were in a feedback loop and their levels were influenced by the genes *LCAT, PLA2G4B*, and *LPCAT* (Supplemental Figure 5.) Previous studies have suggested that *LCAT* and *LPCAT* are related to AD [51,52]. Thus, by connecting glycerophospholipids, especially these two metabolites with t-tau and p-tau, we provide further evidence for their connections with AD pathogenesis.

Our sample sizes were relatively small for both IMPACT and WRAP; however, the 38 CSF metabolites’ associations with CSF t-tau and p-tau levels identified before [12] were replicated in the independent IMPACT data, strengthening our confidence that these 38 metabolites are important for tau pathology. However, further research is necessary to understand whether a causal relationship exists between these CSF metabolites and tau pathology. One limitation of this study is that both IMPACT and WRAP are predominantly non-Hispanic white/Caucasian, so the findings of this study may not be generalizable to other races/ethnicities. Another limitation is that most of the 38 metabolites are highly correlated with each other. LASSO selected seven metabolites that have non-zero effects on tau, but the resulting metabolites are still correlated with each other (Supplemental Figure 6; range of 0.40 to 0.96). A more sophisticated approach that can further remove non-independent metabolites is needed for clinical application. A third limitation is that in our pathway analysis, only three or two metabolites were included in the enriched pathways (pentose and glucoronate interconversions and glycerophospholipid metabolism, respectively). Future research will be necessary to confirm these results.

In summary, we aimed to replicate earlier findings of 38 CSF metabolites’ correlation with tau and expand the biological knowledge of them to better understand their roles in AD pathogenesis. 38 CSF metabolites individually associated with two tau outcomes significantly and, together, explained a large amount of variance in tau. A subset of these metabolites, selected by LASSO, improved the prediction accuracy of AD/MCI vs. HOC over a model that included established predictors of AD. Two promising metabolic pathways, pentose and glucuronate interconversions metabolism and glycerophospholipid metabolism, were identified in this study and have been shown to be related to AD in previous literature. IMPACT and WRAP are ongoing longitudinal studies that are continuing to collect plasma and CSF from study participants, and additional data will be generated in the future. These data may help fill in gaps regarding the mechanisms linking metabolites and AD, improve the establishment of CSF-based metabolite biomarkers, and identify novel drug targets.

## Acknowledgements

The authors especially thank the WRAP and Wisconsin ADRC participants and staff for their contributions to the studies. Without their efforts this research would not be possible. This study was supported by the National Institutes of Health (NIH) grants [R01AG27161 (Wisconsin Registry for Alzheimer Prevention: Biomarkers of Preclinical AD), R01AG054047 (Genomic and Metabolomic Data Integration in a Longitudinal Cohort at Risk for Alzheimer’s Disease), and P30AG062715 (Wisconsin Alzheimer’s Disease Research Center Grant)], the Helen Bader Foundation, Northwestern Mutual Foundation, Extendicare Foundation, State of Wisconsin, the Clinical and Translational Science Award (CTSA) program through the NIH National Center for Advancing Translational Sciences (NCATS) grant [UL1TR000427], and the University of Wisconsin-Madison Office of the Vice Chancellor for Research and Graduate Education with funding from the Wisconsin Alumni Research Foundation. This research was supported in part by the Intramural Research Program of the National Institute on Aging. Computational resources were supported by core grants to the Center for Demography and Ecology [P2CHD047873] and the Center for Demography of Health and Aging [P30AG017266]. Author Y Deming was supported by a training grant from the National Institute on Aging [T32AG000213]. HZ is a Wallenberg Scholar supported by grants from the Swedish Research Council [#2018-02532], the European Research Council [#681712], the Swedish state under the agreement between the Swedish government and the County Councils, the ALF-agreement [#ALFGBG-720931], the Alzheimer Drug Discovery Foundation (ADDF), USA [#201809-2016862], and the UK Dementia Research Institute at UCL.

KB is supported by the Swedish Research Council (#2017-00915), ADDF, USA [#RDAPB-201809-2016615], the Swedish Alzheimer Foundation [#AF-742881], Hjärnfonden, Sweden [#FO2017-0243], the Swedish state under the agreement between the Swedish government and the County Councils, the ALF-agreement [#ALFGBG-715986], and European Union Joint Program for Neurodegenerative Disorders [JPND2019-466-236].

