## Supplemental files for "CSF metabolites associate with CSF tau and improve prediction of Alzheimer’s disease status"

**Supplemental Table 1. Information of the 38 CSF metabolites.**

| **Compound ID** | **Biochemical Names** | **Super pathway** | **Sub-pathway** | **HMDB ID** | **Missing Percentage*** |
| --- | --- | --- | --- | --- | --- |
| 63 | Cholesterol | Lipid | Sterol | HMDB00067 | 0.00% |
| 575 | Arabinose | Carbohydrate | Pentose Metabolism | HMDB00646 | 0.00% |
| 1591 | N-acetylvaline | Amino Acid | Leucine, Isoleucine and Valine Metabolism | HMDB11757 | 0.00% |
| 19130 | 1,2-dipalmitoyl-GPC (16:0/16:0) | Lipid | Phosphatidylcholine (PC) | HMDB00564 | 0.00% |
| 19258 | 1-myristoyl-2-palmitoyl-GPC (14:0/16:0) | Lipid | Phosphatidylcholine (PC) | HMDB07869 | 0.00% |
| 19503 | Stearoyl sphingomyelin (d18:1/18:0) | Lipid | Sphingolipid Metabolism | HMDB01348 | 0.00% |
| 20699 | Erythritol | Xenobiotics | Food Component/Plant | HMDB02994 | 0.27% |
| 27731 | Ribonate | Carbohydrate | Pentose Metabolism | HMDB00867 | 0.00% |
| 32377 | N-acetylneuraminate | Carbohydrate | Aminosugar Metabolism | HMDB00230 | 0.00% |
| 33939 | N-acetylthreonine | Amino Acid | Glycine, Serine and Threonine Metabolism | HMDB62557 | 0.00% |
| 33955 | 1-palmitoyl-GPC (16:0) | Lipid | Lysophospholipid | HMDB10382 | 0.00% |
| 37058 | Succinylcarnitine (C4-DC) | Energy | TCA Cycle | HMDB61717 | 0.00% |
| 37076 | N-acetylserine | Amino Acid | Glycine, Serine and Threonine Metabolism | HMDB02931 | 0.00% |
| 37506 | Palmitoyl sphingomyelin (d18:1/16:0) | Lipid | Sphingolipid Metabolism | HMDB61712 | 0.00% |
| 37529 | Sphingomyelin (d18:1/18:1, d18:2/18:0) | Lipid | Sphingolipid Metabolism | HMDB12101 | 0.00% |
| 42420 | Erythronate | Carbohydrate | Aminosugar Metabolism | HMDB00613 | 0.00% |
| 42463 | Sphingomyelin (d18:1/14:0, d16:1/16:0) | Lipid | Sphingolipid Metabolism | HMDB12097 | 8.70% |
| 46957 | Gulonate | Cofactors and Vitamins | Ascorbate and Aldarate Metabolism | HMDB03290 | 0.54% |
| 46961 | X - 21785 | Unknown | Unknown | Unavailable | 0.00% |
| 47301 | X - 18887 | Unknown | Unknown | Unavailable | 8.70% |
| 47581 | X - 10457 | Unknown | Unknown | Unavailable | 0.00% |
| 47955 | X - 12906 | Unknown | Unknown | Unavailable | 0.00% |
| 48130 | N6-succinyladenosine | Nucleotide | Purine Metabolism, Adenine containing | HMDB00912 | 0.00% |
| 48258 | 1-oleoyl-GPC (18:1) | Lipid | Lysophospholipid | HMDB02815 | 0.00% |
| 48782 | C-glycosyl tryptophan | Amino Acid | Tryptophan Metabolism | Unavailable | 0.00% |
| 48885 | Arabitol/xylitol | Carbohydrate | Pentose Metabolism | HMDB0002917 | 0.00% |
| 49637 | X - 23739 | Unknown | Unknown | Unavailable | 0.82% |
| 52265 | X - 24228 | Unknown | Unknown | Unavailable | 0.00% |
| 52438 | 1-stearoyl-2-oleoyl-GPC (18:0/18:1) | Lipid | Phosphatidylcholine (PC) | HMDB08038 | 0.27% |
| 52461 | 1-palmitoyl-2-oleoyl-GPC (16:0/18:1) | Lipid | Phosphatidylcholine (PC) | HMDB07972 | 0.00% |
| 52470 | 1-palmitoyl-2-palmitoleoyl-GPC (16:0/16:1) | Lipid | Phosphatidylcholine (PC) | HMDB07969 | 0.00% |
| 52525 | X - 24329 | Unknown | Unknown | Unavailable | 0.00% |
| 52616 | 1-palmitoyl-2-stearoyl-GPC (16:0/18:0) | Lipid | Phosphatidylcholine (PC) | HMDB07970 | 1.36% |
| 52769 | X - 24452 | Unknown | Unknown | Unavailable | 0.00% |
| 53127 | X - 24699 | Unknown | Unknown | Unavailable | 0.27% |
| 54923 | Beta-citrylglutamate | Amino Acid | Glutamate Metabolism | Unavailable | 0.00% |
| 57747 | 3-methylglutaconate | Amino Acid | Leucine, Isoleucine and Valine Metabolism | HMDB00522 | 0.00% |
| 62309 | N-acetyl-isoputreanine | Amino Acid | Polyamine Metabolism | Unavailable | 0.00% |

* The missing percentage was calculated based on the combined data set of 368 samples from the IMPACT and WRAP cohorts.

**Supplemental Table 2. Estimates and p-values for 38 individual metabolite-tau associations in the IMPACT.**

|  | **T-tau** | | | **P-tau** | | |
| --- | --- | --- | --- | --- | --- | --- |
| **Biochemical Name** | **Estimate** | **P** | **Adjusted P** | **Estimate** | **P** | **Adjusted P** |
| 1,2-dipalmitoyl-GPC (16:0/16:0) | 659.64 | 2.26E-15 | 8.57E-14 | 71.64 | 5.41E-19 | 2.06E-17 |
| 1-myristoyl-2-palmitoyl-GPC (14:0/16:0) | 547.92 | 8.19E-11 | 3.11E-09 | 61.82 | 3.70E-14 | 1.41E-12 |
| 1-oleoyl-GPC (18:1) | 535.92 | 2.34E-13 | 8.89E-12 | 57.09 | 1.24E-15 | 4.69E-14 |
| 1-palmitoyl-2-oleoyl-GPC (16:0/18:1) | 614.28 | 7.88E-12 | 3.00E-10 | 69.89 | 6.57E-16 | 2.50E-14 |
| 1-palmitoyl-2-palmitoleoyl-GPC (16:0/16:1) | 521.72 | 4.85E-08 | 1.84E-06 | 62.50 | 1.36E-11 | 5.17E-10 |
| 1-palmitoyl-2-stearoyl-GPC (16:0/18:0) | 137.43 | 0.0013596 | 0.0516648 | 17.79 | 2.28E-05 | 8.67E-04 |
| 1-palmitoyl-GPC (16:0) | 533.94 | 3.95E-13 | 1.50E-11 | 57.01 | 1.97E-15 | 7.50E-14 |
| 1-stearoyl-2-oleoyl-GPC (18:0/18:1) | 434.28 | 4.64E-08 | 1.76E-06 | 48.38 | 4.69E-10 | 1.78E-08 |
| 3-methylglutaconate | 619.03 | 8.46E-08 | 3.22E-06 | 72.15 | 1.32E-10 | 5.00E-09 |
| Arabinose | 771.57 | 1.22E-11 | 4.62E-10 | 82.00 | 1.83E-13 | 6.96E-12 |
| Arabitol/xylitol | 687.59 | 3.31E-17 | 1.26E-15 | 73.14 | 2.26E-20 | 8.59E-19 |
| Beta-citrylglutamate | 632.86 | 9.34E-18 | 3.55E-16 | 65.79 | 6.65E-20 | 2.53E-18 |
| C-glycosyltryptophan | 742.27 | 5.40E-22 | 2.05E-20 | 81.58 | 3.86E-29 | 1.47E-27 |
| Cholesterol | 603.22 | 1.83E-11 | 6.94E-10 | 67.08 | 1.30E-14 | 4.92E-13 |
| Erythritol | 328.54 | 6.90E-05 | 0.0026207 | 32.56 | 6.93E-05 | 0.0026325 |
| Erythronate | 1055.07 | 2.62E-14 | 9.96E-13 | 115.27 | 7.93E-18 | 3.02E-16 |
| Gulonate | 268.32 | 1.61E-06 | 6.11E-05 | 30.57 | 2.31E-08 | 8.79E-07 |
| N6-succinyladenosine | 467.77 | 7.07E-12 | 2.69E-10 | 50.83 | 1.97E-14 | 7.50E-13 |
| N-acetyl-isoputreanine | 671.03 | 3.92E-10 | 1.49E-08 | 68.74 | 8.13E-11 | 3.09E-09 |
| N-acetylneuraminate | 672.41 | 5.80E-20 | 2.20E-18 | 72.03 | 2.41E-24 | 9.15E-23 |
| N-acetylserine | 751.01 | 2.53E-12 | 9.60E-11 | 77.87 | 1.50E-13 | 5.68E-12 |
| N-acetylthreonine | 769.28 | 4.04E-18 | 1.54E-16 | 76.66 | 2.37E-18 | 8.99E-17 |
| N-acetylvaline | 652.34 | 8.69E-13 | 3.30E-11 | 69.26 | 7.98E-15 | 3.03E-13 |
| Palmitoyl sphingomyelin (d18:1/16:0) | 531.68 | 1.86E-10 | 7.07E-09 | 57.29 | 2.19E-12 | 8.31E-11 |
| Ribonate | 439.61 | 1.95E-08 | 7.40E-07 | 50.47 | 3.28E-11 | 1.25E-09 |
| Sphingomyelin (d18:1/14:0, d16:1/16:0) | 191.19 | 4.99E-06 | 1.90E-04 | 19.70 | 1.93E-06 | 7.34E-05 |
| Sphingomyelin (d18:1/18:1, d18:2/18:0) | 528.86 | 1.49E-10 | 5.68E-09 | 60.71 | 2.60E-14 | 9.88E-13 |
| Stearoyl sphingomyelin (d18:1/18:0) | 613.62 | 1.83E-14 | 6.95E-13 | 68.38 | 6.57E-19 | 2.50E-17 |
| Succinylcarnitine (C4) | 487.72 | 1.20E-07 | 4.55E-06 | 51.38 | 1.49E-08 | 5.67E-07 |
| X - 10457 | 639.09 | 3.85E-22 | 1.46E-20 | 68.79 | 1.19E-27 | 4.54E-26 |
| X - 12906 | 537.04 | 8.02E-09 | 3.05E-07 | 56.01 | 1.01E-09 | 3.83E-08 |
| X - 18887 | 176.83 | 5.80E-05 | 0.0022025 | 18.96 | 1.24E-05 | 4.72E-04 |
| X - 21785 | 718.75 | 6.65E-10 | 2.53E-08 | 76.36 | 2.30E-11 | 8.73E-10 |
| X - 23739 | 407.90 | 2.70E-08 | 1.02E-06 | 40.61 | 2.30E-08 | 8.74E-07 |
| X - 24228 | 827.46 | 1.90E-23 | 7.21E-22 | 87.28 | 8.50E-28 | 3.23E-26 |
| X - 24329 | 626.37 | 3.17E-14 | 1.20E-12 | 68.03 | 1.78E-17 | 6.77E-16 |
| X - 24699 | 714.07 | 1.02E-21 | 3.86E-20 | 75.45 | 1.05E-25 | 4.00E-24 |
| X - 52769 | 773.74 | 4.97E-16 | 1.89E-14 | 86.03 | 4.60E-21 | 1.75E-19 |

**Supplemental Table 3. The meta-analysis results of 38 individual metabolite-tau associations by WRAP and IMPACT.**

|  | **T-tau** | | | **P-tau** | | |
| --- | --- | --- | --- | --- | --- | --- |
| **Biochemical Name** | **Estimate** | **P** | **Adjusted P** | **Estimate** | **P** | **adjusted P** |
| 1,2-dipalmitoyl-GPC (16:0/16:0) | 426.51 | 1.75E-24 | 5.26E-23 | 63.26 | 2.53E-22 | 7.58E-21 |
| 1-myristoyl-2-palmitoyl-GPC (14:0/16:0) | 335.05 | 3.45E-16 | 1.04E-14 | 52.52 | 3.18E-16 | 9.54E-15 |
| 1-oleoyl-GPC (18:1) | 257.16 | 3.04E-16 | 9.11E-15 | 33.26 | 4.74E-10 | 1.42E-08 |
| 1-palmitoyl-2-oleoyl-GPC (16:0/18:1) | 315.12 | 7.44E-15 | 2.23E-13 | 53.64 | 1.03E-14 | 3.10E-13 |
| 1-palmitoyl-2-palmitoleoyl-GPC (16:0/16:1) | 318.86 | 7.90E-13 | 2.37E-11 | 53.83 | 2.98E-13 | 8.93E-12 |
| 1-palmitoyl-2-stearoyl-GPC (16:0/18:0) | 104.53 | 1.43E-06 | 4.28E-05 | 22.56 | 8.58E-08 | 2.57E-06 |
| 1-palmitoyl-GPC (16:0) | 224.23 | 3.11E-14 | 9.32E-13 | 29.21 | 9.93E-09 | 2.98E-07 |
| 1-stearoyl-2-oleoyl-GPC (18:0/18:1) | 209.85 | 2.01E-10 | 6.02E-09 | 35.95 | 3.96E-09 | 1.19E-07 |
| 3-(3-amino-3-carboxypropyl)uridine | 264.81 | 1.48E-14 | 4.43E-13 | 42.00 | 1.55E-12 | 4.64E-11 |
| 3-methylglutaconate | 540.68 | 4.66E-19 | 1.40E-17 | 79.00 | 4.14E-19 | 1.24E-17 |
| Arabinose | 493.56 | 4.94E-16 | 1.48E-14 | 72.31 | 7.42E-12 | 2.23E-10 |
| Arabitol/xylitol | 615.10 | 4.71E-44 | 1.41E-42 | 81.71 | 3.46E-36 | 1.04E-34 |
| Beta-citrylglutamate | 288.90 | 1.41E-19 | 4.24E-18 | 39.71 | 8.54E-13 | 2.56E-11 |
| C-glycosyltryptophan | 550.97 | 6.11E-45 | 1.83E-43 | 72.51 | 8.58E-35 | 2.57E-33 |
| Cholesterol | 213.54 | 1.29E-11 | 3.88E-10 | 27.41 | 6.88E-07 | 2.06E-05 |
| Erythritol | 13.25 | 2.07E-01 | 1.00 | 2.99 | 1.38E-01 | 1.00 |
| Erythronate | 533.29 | 2.36E-15 | 7.09E-14 | 94.97 | 3.02E-17 | 9.07E-16 |
| Gulonate | 285.94 | 2.71E-15 | 8.13E-14 | 55.42 | 3.18E-18 | 9.54E-17 |
| Hydroxy-N6,N6,N6-trimethyllysine | 561.79 | 2.22E-28 | 6.65E-27 | 70.54 | 3.13E-18 | 9.39E-17 |
| N6-succinyladenosine | 405.08 | 7.13E-26 | 2.14E-24 | 62.75 | 1.77E-25 | 5.30E-24 |
| N-acetyl-isoputreanine | 506.94 | 2.21E-18 | 6.63E-17 | 63.88 | 4.87E-13 | 1.46E-11 |
| N-acetylneuraminate | 552.51 | 2.47E-48 | 7.41E-47 | 69.00 | 1.23E-40 | 3.69E-39 |
| N-acetylserine | 563.79 | 4.89E-24 | 1.47E-22 | 78.78 | 4.61E-21 | 1.38E-19 |
| N-acetylthreonine | 320.06 | 4.75E-17 | 1.42E-15 | 54.85 | 1.58E-16 | 4.75E-15 |
| N-acetylvaline | 286.62 | 4.02E-14 | 1.21E-12 | 44.53 | 4.59E-11 | 1.38E-09 |
| Palmitoyl sphingomyelin (d18:1/16:0) | 277.12 | 4.54E-13 | 1.36E-11 | 37.22 | 1.89E-08 | 5.67E-07 |
| Ribonate | 196.11 | 2.38E-10 | 7.15E-09 | 32.05 | 7.57E-09 | 2.27E-07 |
| Sphingomyelin (d18:1/14:0, d16:1/16:0) | 74.80 | 3.03E-05 | 9.09E-04 | 12.94 | 5.21E-05 | 1.56E-03 |
| Sphingomyelin (d18:1/18:1, d18:2/18:0) | 263.78 | 8.37E-13 | 2.51E-11 | 39.98 | 1.75E-10 | 5.24E-09 |
| Stearoyl sphingomyelin (d18:1/18:0) | 414.50 | 7.34E-24 | 2.20E-22 | 57.59 | 5.34E-18 | 1.60E-16 |
| Succinylcarnitine (C4) | 371.20 | 2.96E-16 | 8.87E-15 | 54.39 | 1.08E-17 | 3.24E-16 |
| X - 10457 | 532.21 | 6.79E-52 | 2.04E-50 | 68.76 | 4.34E-41 | 1.30E-39 |
| X - 12906 | 162.12 | 1.37E-05 | 4.11E-04 | 32.09 | 2.70E-06 | 8.10E-05 |
| X - 18887 | 60.98 | 2.65E-06 | 7.96E-05 | 9.17 | 1.59E-04 | 4.76E-03 |
| X - 21785 | 273.60 | 3.37E-08 | 1.01E-06 | 65.96 | 4.59E-15 | 1.38E-13 |
| X - 23739 | 106.98 | 5.40E-06 | 1.62E-04 | 18.35 | 6.28E-05 | 1.88E-03 |
| X - 24228 | 608.98 | 3.02E-46 | 9.05E-45 | 77.56 | 7.07E-32 | 2.12E-30 |
| X - 24699 | 389.77 | 2.49E-31 | 7.46E-30 | 57.98 | 1.92E-26 | 5.76E-25 |

**Supplemental Table 4. Prediction of 34 plasma metabolites and the same 34 CSF metabolites.**

|  | **IMPACT** | | **WRAP** | |
| --- | --- | --- | --- | --- |
|  | **T-tau** | **P-tau** | **T-tau** | **P-tau** |
| **Demographics and 38 CSF metabolites using baseline samples** | NA^*^ | | 0.634 | 0.688 |
| **Demographics and 34 Plasma metabolites** | NA^*^ | | 0.269 | 0.301 |
| **Demographics and 34 Plasma metabolites using baseline samples** | NA^*^ | | 0.260 | 0.272 |
| **Demographics and 34 CSF metabolites** | 0.725 | 0.784 | 0.621 | 0.641 |

^*^ The IMPACT used cross-sectional samples and the plasma data was not available in IMPACT.

**Supplemental Table 5.** **Factor loadings for 38 CSF metabolites in the IMPACT and WRAP.**

| **Biochemical Name** | **Super-pathway** | **Sub-pathway** | **IMPACT** | | | **WRAP** | | |
| --- | --- | --- | --- | --- | --- | --- | --- | --- |
|  |  |  | **Factor 1** | **Factor 2** | **Factor 3** | **Factor 1** | **Factor 2** | **Factor 3** |
| **X - 24699** | **Unknown** | **Unknown** | **0.917** | 0.022 | -0.002 | **0.800** | 0.093 | -0.015 |
| **X - 24452** | **Unknown** | **Unknown** | **0.855** | -0.083 | 0.210 | **0.749** | -0.027 | 0.289 |
| **N-acetylserine** | **Amino Acid** | **Glycine, Serine and Threonine Metabolism** | **0.850** | 0.048 | -0.292 | **0.924** | -0.037 | -0.168 |
| **N-acetylthreonine** | **Amino Acid** | **Glycine, Serine and Threonine Metabolism** | **0.833** | -0.030 | 0.126 | **0.863** | -0.041 | 0.129 |
| **Erythronate** | **Carbohydrate** | **Aminosugar Metabolism** | **0.826** | 0.014 | -0.167 | **0.817** | 0.061 | -0.064 |
| **X - 10457** | **Unknown** | **Unknown** | **0.826** | 0.030 | 0.143 | **0.700** | 0.179 | 0.086 |
| **N-acetyl-isoputreanine** | **Amino Acid** | **Polyamine Metabolism** | **0.822** | -0.102 | 0.118 | **0.794** | -0.075 | 0.062 |
| **C-glycosyl tryptophan** | **Amino Acid** | **Tryptophan Metabolism** | **0.806** | 0.021 | 0.284 | **0.714** | 0.083 | 0.258 |
| **Arabitol/xylitol** | **Carbohydrate** | **Pentose Metabolism** | **0.798** | 0.120 | -0.123 | **0.777** | 0.143 | 0.013 |
| **X - 24228** | **Unknown** | **Unknown** | **0.783** | 0.081 | 0.169 | **0.699** | 0.181 | 0.070 |
| **3-methylglutaconate** | **Amino Acid** | **Leucine, Isoleucine and Valine Metabolism** | **0.775** | 0.034 | -0.278 | **0.811** | 0.064 | -0.156 |
| **Ribonate** | **Carbohydrate** | **Pentose Metabolism** | **0.763** | -0.017 | -0.056 | **0.833** | -0.056 | -0.074 |
| **N-acetylvaline** | **Amino Acid** | **Leucine, Isoleucine and Valine Metabolism** | **0.739** | 0.016 | 0.091 | **0.883** | -0.258 | 0.159 |
| **X - 24329** | **Unknown** | **Unknown** | **0.732** | 0.064 | 0.117 | **0.819** | -0.057 | 0.047 |
| **N-acetylneuraminate** | **Carbohydrate** | **Aminosugar Metabolism** | **0.715** | 0.250 | -0.248 | **0.762** | 0.208 | -0.189 |
| **Succinylcarnitine (C4-DC)** | **Energy** | **TCA Cycle** | **0.689** | -0.002 | 0.035 | **0.676** | 0.074 | 0.121 |
| **Arabinose** | **Carbohydrate** | **Pentose Metabolism** | **0.683** | 0.128 | -0.205 | **0.641** | 0.090 | 0.025 |
| **Beta-citrylglutamate** | **Amino Acid** | **Glutamate Metabolism** | **0.678** | 0.077 | 0.185 | **0.668** | 0.069 | 0.211 |
| **N6-succinyladenosine** | **Nucleotide** | **Purine Metabolism, Adenine containing** | **0.644** | 0.180 | -0.110 | **0.729** | 0.139 | 0.070 |
| **X - 12906** | **Unknown** | **Unknown** | **0.642** | 0.026 | 0.130 | **0.648** | 0.075 | -0.065 |
| **X - 21785** | **Unknown** | **Unknown** | **0.632** | 0.063 | -0.052 | **0.526** | 0.337 | -0.099 |
| **Erythritol** | **Xenobiotics** | **Food Component/Plant** | **0.542** | -0.092 | -0.158 | **0.327** | 0.077 | -0.071 |
| **X - 18887** | **Unknown** | **Unknown** | **0.469** | 0.049 | 0.015 | **0.398** | 0.116 | 0.098 |
| **X - 23739** | **Unknown** | **Unknown** | **0.378** | 0.262 | 0.098 | **0.450** | 0.164 | 0.155 |
| **Gulonate** | **Cofactors and Vitamins** | **Ascorbate and Aldarate Metabolism** | **0.327** | 0.364 | -0.284 | **0.678** | 0.215 | -0.177 |
| **Stearoyl sphingomyelin (d18:1/18:0)** | **Lipid** | **Sphingolipid Metabolism** | 0.204 | **0.789** | 0.038 | 0.209 | **0.736** | 0.124 |
| **1,2-dipalmitoyl-GPC (16:0/16:0)** | **Lipid** | **Phosphatidylcholine (PC)** | 0.170 | **0.846** | -0.095 | 0.283 | **0.783** | -0.111 |
| **Cholesterol** | **Lipid** | **Sterol** | 0.099 | **0.726** | 0.146 | 0.228 | **0.453** | 0.200 |
| **1-palmitoyl-2-oleoyl-GPC (16:0/18:1)** | **Lipid** | **Phosphatidylcholine (PC)** | 0.095 | **0.918** | -0.080 | 0.197 | **0.845** | -0.074 |
| **1-palmitoyl-2-stearoyl-GPC (16:0/18:0)** | **Lipid** | **Phosphatidylcholine (PC)** | 0.062 | **0.635** | -0.120 | 0.206 | **0.618** | 0.108 |
| **Sphingomyelin (d18:1/18:1, d18:2/18:0)** | **Lipid** | **Sphingolipid Metabolism** | 0.046 | **0.862** | 0.141 | -0.068 | **0.889** | 0.146 |
| **1-stearoyl-2-oleoyl-GPC (18:0/18:1)** | **Lipid** | **Phosphatidylcholine (PC)** | 0.013 | **0.841** | -0.018 | 0.131 | **0.838** | 0.017 |
| **1-myristoyl-2-palmitoyl-GPC (14:0/16:0)** | **Lipid** | **Phosphatidylcholine (PC)** | -0.043 | **0.956** | -0.090 | -0.020 | **0.934** | -0.074 |
| **Sphingomyelin (d18:1/14:0, d16:1/16:0)** | **Lipid** | **Sphingolipid Metabolism** | -0.072 | **0.680** | 0.186 | -0.120 | **0.706** | 0.272 |
| **Palmitoyl sphingomyelin (d18:1/16:0)** | **Lipid** | **Sphingolipid Metabolism** | -0.093 | **0.953** | 0.150 | -0.124 | **0.902** | 0.181 |
| **1-palmitoyl-2-palmitoleoyl-GPC (16:0/16:1)** | **Lipid** | **Phosphatidylcholine (PC)** | -0.100 | **1.004** | -0.062 | 0.015 | **0.920** | -0.057 |
| **1-oleoyl-GPC (18:1)** | **Lipid** | **Lysophospholipid** | 0.335 | 0.348 | **0.560** | 0.182 | 0.158 | **0.760** |
| **1-palmitoyl-GPC (16:0)** | **Lipid** | **Lysophospholipid** | 0.266 | 0.421 | **0.551** | 0.095 | 0.223 | **0.770** |
|  |  | **Proportion Variance** | 0.352 | 0.198 | 0.049 | 0.353 | 0.225 | 0.039 |
|  |  | **Cumulative Variance** | 0.352 | 0.551 | 0.600 | 0.353 | 0.577 | 0.616 |

Orange cells denote factor 1, green cells denote factor 2, and blue cells denote factor 3.

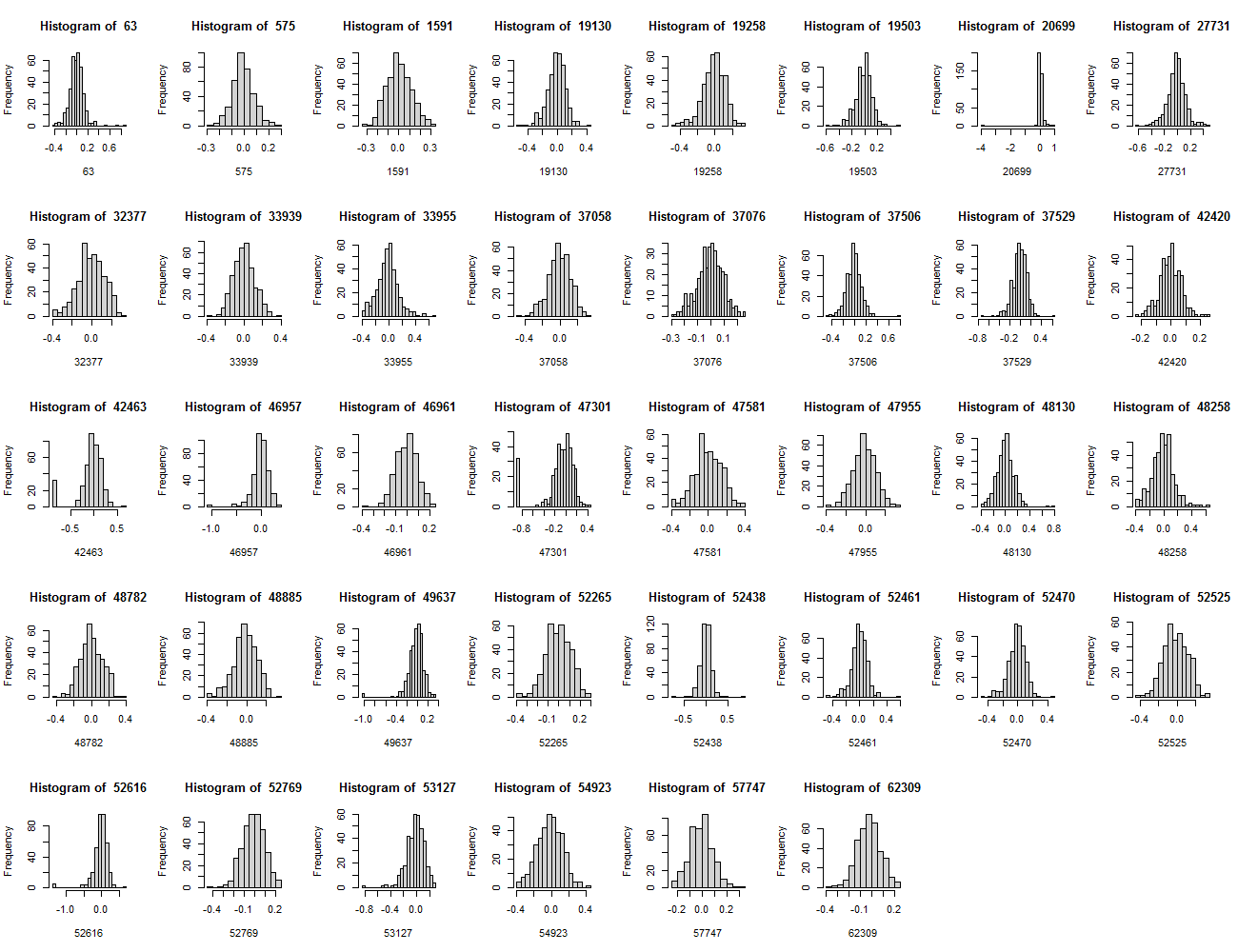

**Supplemental Figure 1. The distributions of 38 CSF metabolites after imputation and Log10 transformation based on** the combined data set of 368 samples from the **IMPACT and WRAP cohorts.**

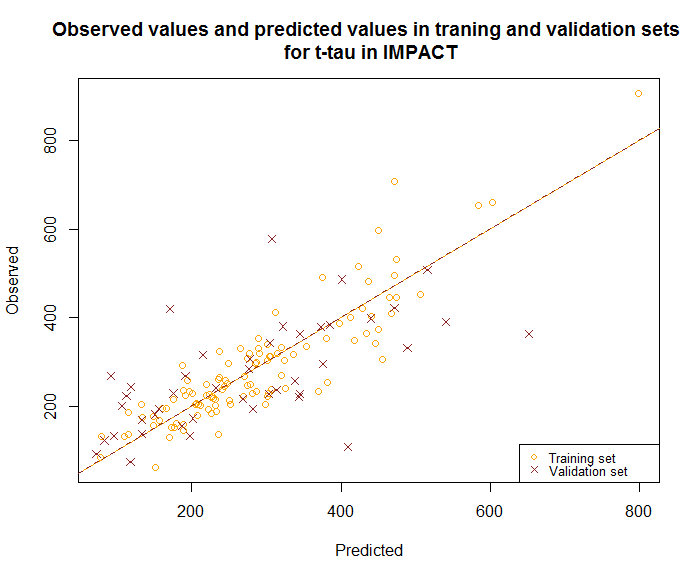

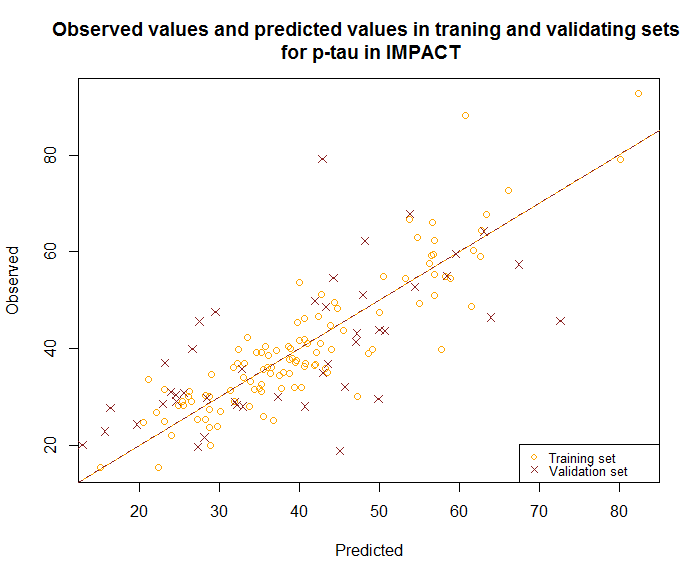

**(a)	(b)**
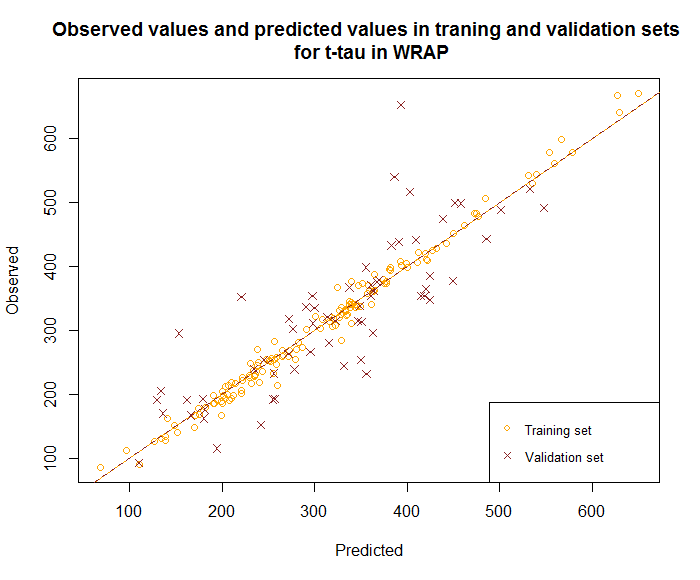

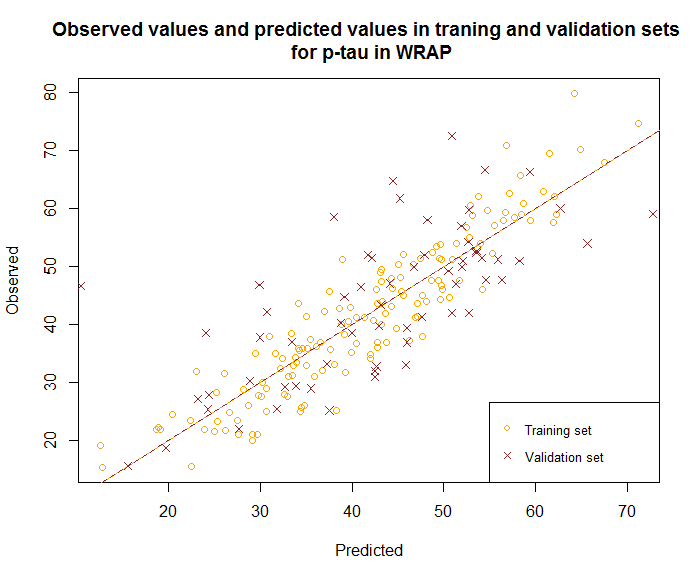

**(c)	(d)**

**Supplemental Figure 2. Scatter plots of observed and predicted values from training and validation sets.** The yellow circles represent the training set values and the brown crosses represent the predicted values. The outcome and cohort from (a) to (d) are (a) t-tau in the IMPACT, (b) p-tau in the IMPACT, (c) t-tau in WRAP, and (d) p-tau in WRAP. The dashed lines are from 0 to 1. If the observed values perfectly equal to predicted values, they will lie on the dash lines.

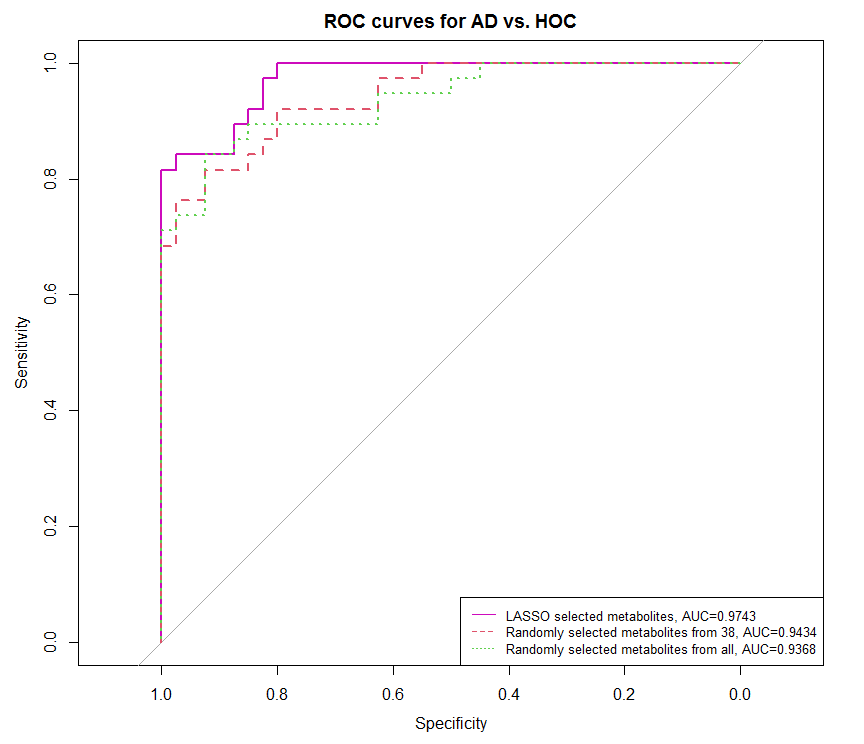

**(a)**

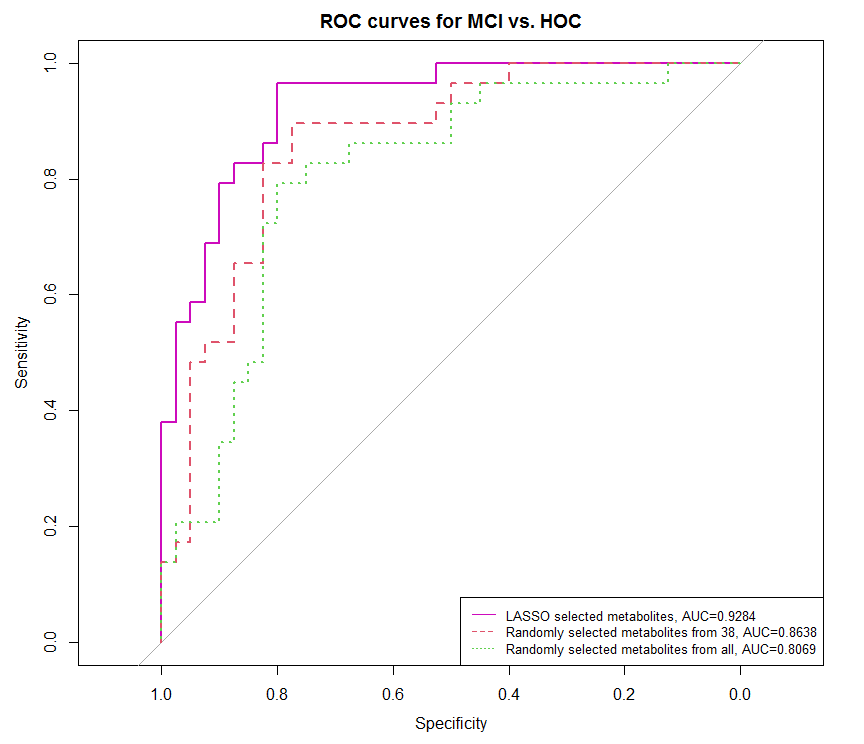

**(b)**

**Supplemental Figure 3. ROC curves and AUC scores of predictions by 6 models in the Wisconsin ADRC (a) AD vs. HOC (b) MCI vs. HOC.** Seven LASSO selected metabolites N-acetylneuraminate, C-glycosyl tryptophan, X-10457, X-24228, 1-oleoyl-GPC(18:1), 1-palmitoyl-GPC(16:0), and 1-myristoyl-2-palmitoyl-GPC(14:0/16:0); seven randomly selected from 38 metabolites are: 1-myristoyl-2-palmitoyl-GPC (14:0/16:0), 1-palmitoyl-2-stearoyl-GPC (16:0/18:0), 1-palmitoyl-2-oleoyl-GPC (16:0/18:1), hydroxy-N6,N6,N6-trimethyllysine, ribonate, X – 18887, and arabitol/xylitol; seven randomly selected metabolites from all metabolites are: tryptophan betaine, 3-hydroxybutyrate (BHBA), pyroglutamine, dimethylglycine, indole acetate, pipecolate, and methyl succinate. All models adjusted for age, sex, years of education, *APOE* ε4 count, t-tau, p-tau, and Aβ42.

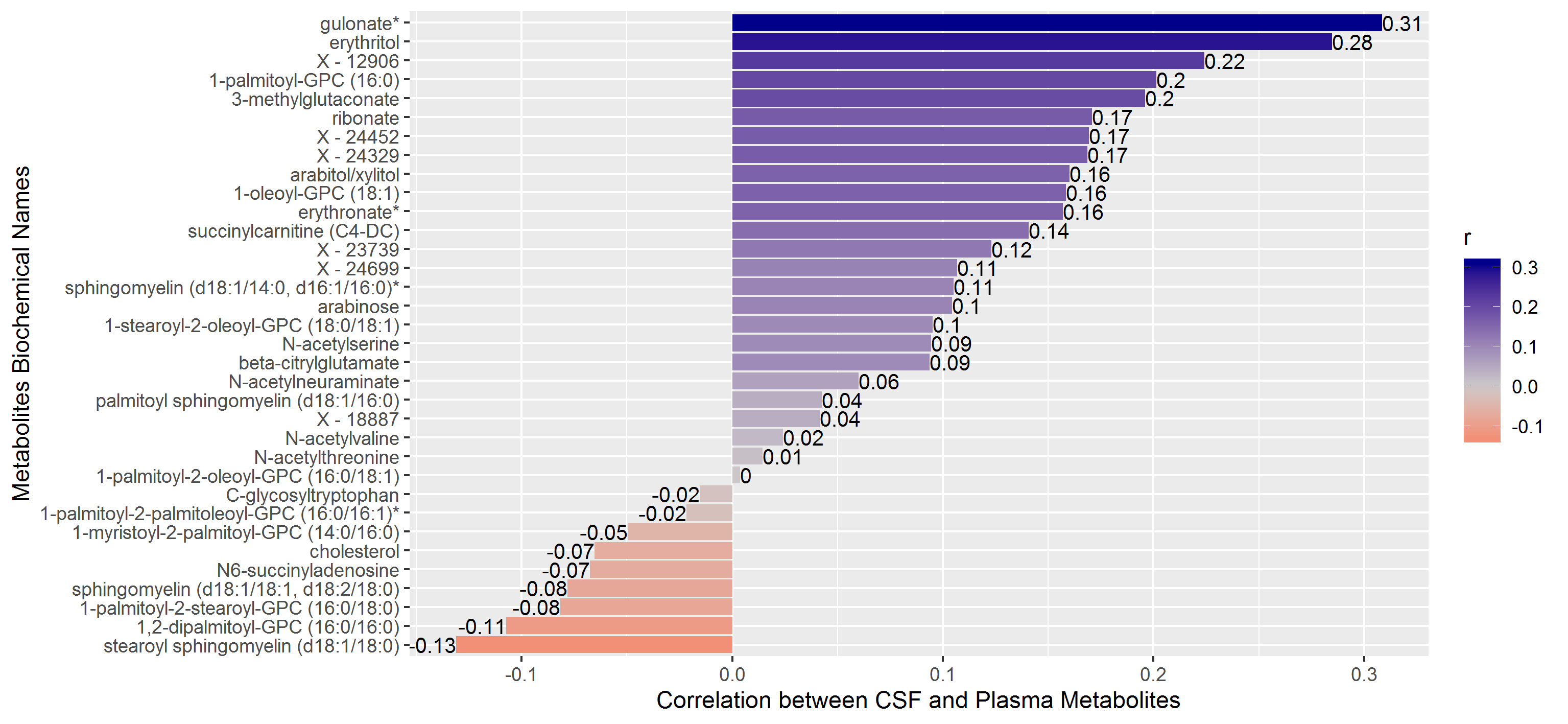

**Supplemental Figure 4. Correlations between 34 metabolites present in both CSF and plasma.** Positive correlations are displayed in blue and negative correlations in red color. The length of the bar is proportional to the correlation coefficients. Here, the correlations range from 0.31 to -0.13 [9].

**1-palmitoyl-2-palmitoleoyl-GPC(16:0/16:1)**

**1-oleoyl-GPC(18:1)**

*LPCAT*

*LCAT*

*PLA2G4B*

**Supplemental Figure 5. The relation between two metabolites in the glycerophospholipid metabolism pathway based on the KEGG pathway database.**

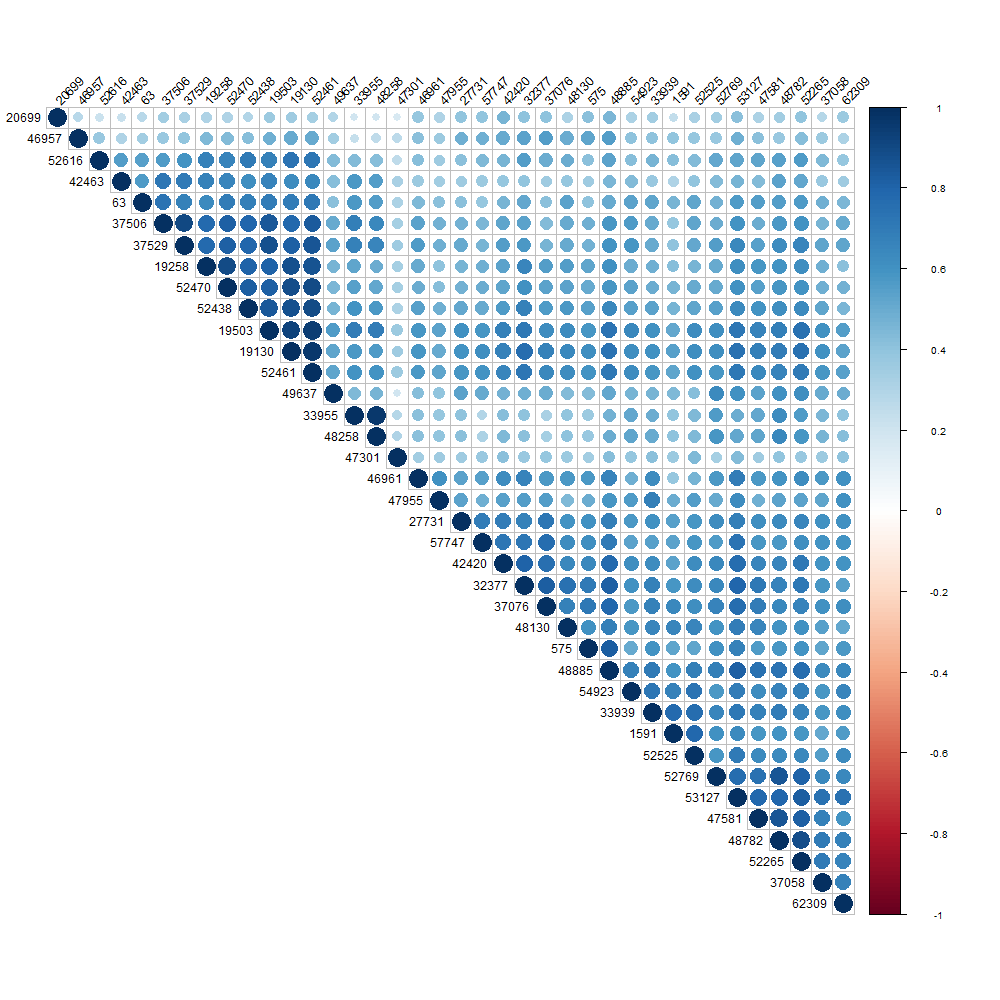

**Supplemental Figure 6. Pairwise correlation of 38 CSF metabolites.** Positive correlations are displayed in blue and negative correlations in red color. Color intensity and the size of the circle are proportional to the correlation coefficients. Here, 38 CSF metabolites are positively correlated with each other, and all correlations are statistically significant. The metabolite compound IDs are shown here. Refer to Supplemental Table 1 for metabolite biochemical names.
